# ViMIC: A Database of Human Disease-related Virus Mutations, Integration Sites and Cis-effects

**DOI:** 10.1101/2020.10.28.359919

**Authors:** Ying Wang, Yuantao Tong, Zeyu Zhang, Rongbin Zheng, Danqi Huang, Jinxuan Yang, Hui Zong, Fanglin Tan, Xiaoyan Zhang

## Abstract

Molecular mechanisms of virus-related diseases involve multiple factors, including viral mutation accumulation and integration of a viral genome into the host DNA. With increasing attention being paid to virus-mediated pathogenesis and the development of many useful technologies to identify virus mutations (VMs) and viral integration sites (VISs), abundant literatures on these topics are available in PubMed. However, knowledge of VMs and VISs is widely scattered in numerous published papers, and the association of VMs with VISs in the viral genome or the functional annotation of VISs still lacks integration and curation. To address these challenges, we built a database of human disease-related **Vi**rus **M**utations, **I**ntegration sites and **C**is-effects (ViMIC), which specialize in three features: virus mutation sites, viral integration sites and target genes. In total, the ViMIC provides information on 6,461 VMs, 79,089 VISs, and 15,056 viral target genes of 8 viruses in 65 human diseases obtained from literatures. Furthermore, in ViMIC, users are allowed to explore the cis-effects of virus-host interactions by surveying 78 histone modifications, binding of 1,358 transcription regulators, and chromatin accessibility on these VISs. We believe ViMIC will become a valuable resource for the virus research community. The database is available at http://bmtongji.cn/ViMIC/index.php.

## INTRODUCTION

Viruses are of substantial interest due to the seriousness of their associated diseases in humans. The mechanism of viral pathogenesis is complicated, as viruses interact with the host genome on multiple levels (1,2). For example, studies revealed that a virus may exert cis-effects by integrating its genome into human chromosomes (3–6). Mutations in viral regions or genes may also contribute to an increased risk for disease progression and drug resistance (7–9). Some virus mutations (VMs) may play an important role in promoting the occurrence of diseases by increasing or decreasing the incidence of viral DNA integration into host cells (10–13). Moreover, alterations in epigenetics may also be induced by the viral protein to promote activation or inhibition of target genes (14–17).

At present, millions of virus studies can be found in PubMed. Such studies have generated vast amounts of information relevant to VMs and viral integration sites (VISs) covering different types of viruses and human diseases. Moreover, domain knowledge that can be used to improve the understanding of virus cis-effects, for instance, gene expression and transcription cis-regulatory information, has been deposited in freely accessible databases, such as Cistrome Data Browser (18,19) and the Gene Expression Omnibus (GEO) (20). However, the well-curated information and data are still distributed in separate public domains that lack integration and standardization. Therefore, it is quite challenging for researchers to quickly search, visualize, and re-use such knowledge for their studies of virus-host interactions. Several databases, such as the Viral Integration Site DataBase (VISDB) (21), Dr.VIS (22), the Retrovirus Integration Database (RID) (23) and the ISDB (24) have been developed. However, to our knowledge, these databases focused on virus integration, and none of existing databases or web-servers are designed for systematically evaluating VMs and functionally annotating of VISs. Understanding of viral pathogenic mutations will abet the effective surveillance and early detection for the high-risk virus-infected diseases. The significant viral mutations or regions for affecting integration reactions have also been reported (10,11,13,25). Besides, previous studies indicated that the transcriptional regulators contribute to the control of viral replication and regulate critical cis-elements in a virus (26,27).

Therefore, to overcome the challenges mentioned above, we present a database of human disease-related **Vi**rus **M**utations, **I**ntegration sites and **C**is-effects (ViMIC). VIMIC is built to fill such gaps from the aspects of virus mutation, integration, and cis-effect of host. The unique of our database over other databases includes: ViMIC is the first database to provide virus mutation information retrieved from a large collection of published literature; ViMIC is the first to integrate VMs with VISs information from the literature; and ViMIC is the first to integrate VIS data with cistrome resources to help researchers who are interested in gene regulation mechanisms to systematically investigate the functional annotation of VISs in the host genome. ViMIC mainly covers three special features, including virus mutation sites, viral integration sites and target genes. First, the database facilitates the search for the VM annotations and the association between VMs and VISs on the viral reference genome. Second, the database can help users determine whether the integration site of viral fragments integrated into the host genome is in a functional element, distinguishing by histone modification, transcription factor binding and chromatin accessibility. Third, the database provides information on host target genes affected by the viral insertion or by the virus gene/protein/region, and on the correlation between gene expression levels and the fraction of infiltrating immune cells in human diseases.

## CONSTRUCTION AND CONTENTS

A schematic overview of the ViMIC data collection, text-mining system, data processing and web interface is summarized in **Figure 1**.

**Figure 1.**
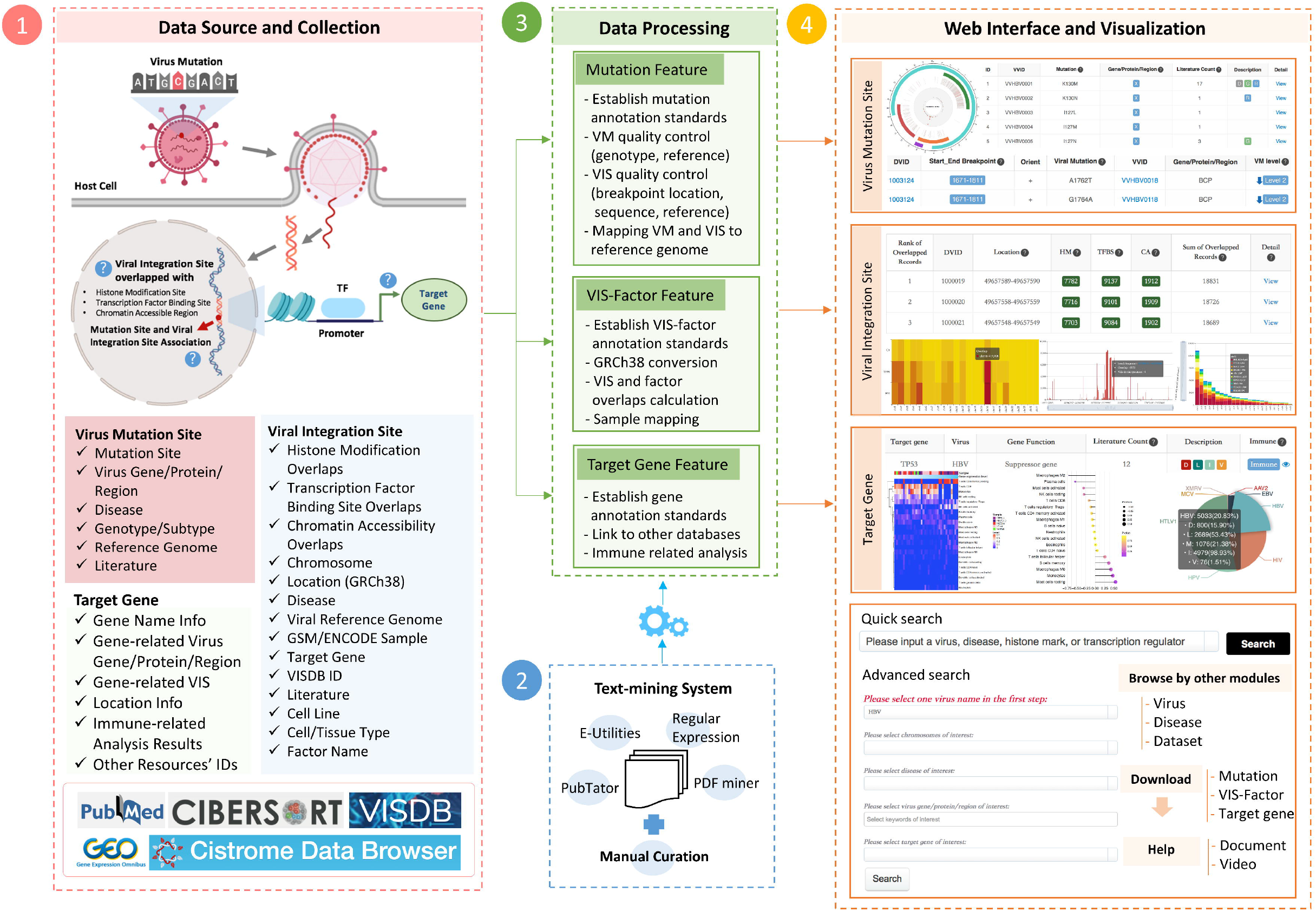
A schematic overview of the ViMIC database. ViMIC collects publicly available virus mutations (VMs), viral integration sites (VISs), cistrome factors and target genes information from multiple public resources, including PubMed, Cistrome Data Browser, VISDB, GEO, etc. ViMIC uses text mining to extract the virus-related bioconcepts from published citations, followed by a manual curation to ensure annotation accuracy. The data processing layer is responsible for specific tasks, such as establishing annotation standards stored in a MySQL relationship database, performing VM-VIS mapping to the reference genome, calculating overlaps between VIS and cistrome factors, analysing the correlation between gene expression and infiltrating immune cell fractions and generating statistical plots. We designed the ViMIC interface with three main features: virus mutation sites, viral integration sites and target genes. ViMIC provides three ways to query and explore data: by keyword search, quick search engine or an advanced search menu. In addition, ViMIC includes other modules, labelled as Virus, Disease, Dataset, Download and Help that allow users to better explore the collected data.

### Data sources

The major sources of data in ViMIC were derived from abundant published literatures, the Cistrome Data Browser (18,19), VISDB (21) and GEO database (20). Viral annotation information was retrieved from a large collection of published literature through our text-mining system, which was followed by a manual curation process. The viral integration sites were retrieved from published literature and VISDB. The human transcription regulator binding, histone modification, and chromatin accessibility data were obtained from the Cistrome Data Browser. Moreover, for each target gene, ViMIC collected gene expression datasets of virus-infected diseases from GEO as well as the Cancer Genome Atlas (TCGA). Next, we conducted a correlation analysis between target gene expressions and fractions of infiltrating immune cells estimated by Cell-type Identification By Estimating Relative Subsets Of RNA Transcripts (CIBERSORT) (28).

### Data retrieval and text mining

We used the Entrez Programming Utilities (E-Utilities) tool to build a set of full names and abbreviations of virus keywords into a fixed URL syntax to search and retrieve virus-related articles from PubMed. A Python script was run to filter the articles by matching keywords (e.g., mutation) as regular expressions. Full-text articles from PMC and abstracts from PubMed (BioC-XML format) with annotations of biomedical concepts in the Bioconepts2pubtatorcentral file were downloaded from PubTatorCentral (29), and this file was then used to extract biomedical entities such as genes, mutations, diseases and chemicals. We also developed a Python script to automate the entity recognition task for those articles without PubTator annotation. The procedure includes literature downloading, conversion of PDF documents to plain text files (PDFMiner library), and noise filtering and information extraction for sentences containing mutations. Mutation entries were automatically extracted from PubMed and PubTator annotation file (30), followed by a manual curation to ensure the data quality of our database.

### VIS quality control in VM-VIS association module

Some of the collected VIS information may be incomplete, which leads to no way to map the VM-VIS association, so we excluded those VISs that did not provide breakpoint locations and the viral reference genome. For each VIS, we define a level index to evaluate the quality of VIS positioning on the viral reference genome.

1. Level 1: The breakpoint locations in the viral reference genome and the sequence of the integrated fragment are both provided. We further used the multiple sequence alignment tool CLUSTALW (31) to check whether the information of this VIS was correct. In indel cases, all quality levels of VISs are Level 1.
2. Level 2: The precise start and end breakpoint locations in the viral reference genome are provided, while the sequence of the viral integration fragment is missing.
3. Level 3: Only one precise breakpoint location in the viral reference genome is provided, and the sequence of the viral integration fragment is missing. In this level, we can only use this breakpoint position in the virus reference genome to infer whether this specific breakpoint is a well-studied VM.

### VM quality control in VM-VIS association module

For each VM and VIS, we retrieved its genotype/subtype and reference sequence of the virus from the literature. As the collected reference genome information between VMs and VISs may not be matched exactly, we also defined a level index to evaluate the quality of the VM positioning on the viral reference genome for each VM.

1. Level 1: The reference genomes of VMs and VISs are exactly matched.
2. Level 2: The virus genotype/subtype of VMs and VISs are exactly matched. However, the reference genomes of VMs and VISs are not matched, but the genomic-length of two reference sequences is consistent.
3. Level 3: The virus genotype/subtype of VMs and VISs are not matched, but the genomic-length of two reference sequences is consistent. For example, HBV mutations, such as K130M, was frequent in both genotype C and B patients (32–34).
4. Level 4: The reference genome of VMs in the literature is missing. In these cases, we only annotated those VMs whose mutant base/amino acid in the wild-type sequence is consistent with the base/amino acid in the same position of the given reference sequence.

All VMs (Level 1-4) which mutant base/amino acid in the wild-type sequence (e.g., G in the G1896A or K in the K130M) must be consistent with the base/amino acid in the same position of the given reference sequence (e.g., the 130^th^ amino acid in the X region is K in the ref. NC_003977).

### VIS-Cistrome Data (Factor) Overlapping Analysis

To construct features for exploring the functional integration of a VIS (VIS-factors), we calculated the overlaps between the VIS and cistromes. The concept of “cistrome” has been widely used in the gene regulation field (18,19,35). As these studies state, a cistrome refers to “the set of cis-acting targets of a trans-acting factor on a genome-wide scale, also known as the in vivo genome-wide location of transcription factor binding-sites or histone modifications.” Therefore, the host gene expression may be influenced if the virus integrates into the Cistrome. Cistrome DB collects human and mouse ChIP-seq, DNase-seq, and ATAC-seq data from publicly available resources. In our current study, we applied Cistrome DB data as cistrome factor information to integrate with data from a virus integration site and assist users to explore the cis-effect of VIS. The procedure included four steps.

First, we derived the viral integration sites in the human genome from VISDB (76,264 curated VISs) and from the PubMed literature (2825 VISs, 11 articles) that reported disease-related viral integration events in the human genome. The collected VISs were processed into BED format, which was then converted into human reference genome coordinates in GRCh38 by the LiftOver if a previous genome version was used. Second, we downloaded the ChIP-seq peaks of human transcription factor and histone modifications as well as the DNase-seq and ATAC-seq peaks from Cistrome Data Browser. We then compressed this information into bgzipped (.gz) files (a broad peak file using all peaks and a narrow peak file using 5-fold-enriched peaks). Third, a GIGGLE (genome search engine) (36) index was built using the compressed peak files, and we used GIGGLE to identify and search the significance of shared genomic loci (overlap) between the VIS and three genome interval files (histone modification site, transcription factor binding site and chromatin accessible region). Finally, we incorporated the GSMID and biological resource information from Cistrome Data Browser into the VIS-cistrome entry (overlap > 0). If a VIS overlapped with cistrome factor-related GSM samples, it would indicate that the virus integration site is a potentially functional DNA element that can influence gene expression in the human.

### Database contents

ViMIC is characterized by a comprehensive curation of VMs, functional annotations of VISs in the host genome and target genes. For each collected virus, ViMIC provides three features, as shown in **Figure 1**.

#### Feature I – Virus Mutation Site

The “Virus Mutation Site” feature provides two modules for users: the VM annotation module and the VM-VIS association module. The VM annotation module includes mutation site, mutation level, virus gene/protein/region, virus-associated disease, genotype/subtype, literature evidence, etc. We designed a description column to indicate more detailed information for each mutation site using the following tags: (i) D: drugs mentioned in the relevant literature; (ii) G: genotypes/subtypes mentioned in the relevant literature; (iii) R: reference genomes mentioned in the relevant literature. In the current version of VM-VIS association module, VISs of 5 viruses including hepatitis B virus (HBV), human papillomavirus virus (HPV), Epstein-Barr virus (EBV), adenovirus associated virus type 2 (AAV2) and Merkel cell polyomavirus (MCV), with precise breakpoint coordinates and the reference genome in the integration literature, were used to map the association between the VMs and VISs.

#### Feature II – Viral Integration Site

The second feature is the VIS-cistrome factor overlap calculation, which we specifically designed to explore whether the VIS is involved in a functional region in the host genome. Therefore, for each VIS, if there are overlaps between the VIS and the genomic peaks of the cistrome factors (sum of overlaps > 0), the detailed results are presented, including the overlap number and annotation of each cistrome sample (GSMID, cell line, cell type, tissue type and factor name). Furthermore, we summarized all the VIS-cistrome factor overlapped records on human chromosomes using heatmap and histogram plots, and the top-ranked cistrome factors ordered by the overlap number in each chromosome using stacked bar plots. Thus, users can view the overlap distribution between the VIS and cistrome factors in the human genome.

#### Feature III – Target gene

This feature provides information for the target genes affected by viral insertion or by the virus gene/protein/region. We have systematically annotated the target gene information, including official gene symbols, official full names, gene aliases, transcripts, gene types, gene location information, gene functions, gene IDs, drug information, etc. For each gene, if there were available gene expression data for patients with the virus-related disease, we further analysed the relationship between its expression and immune cell infiltration and represented it by heatmap and lollipop plots.

In the current version (as of August 31, 2020), the database contains 8 viruses, including HBV, HPV, EBV, AAV2, MCV, human immunodeficiency virus (HIV), human T-cell lymphotropic virus type-1 (HTLV1) and xenotropic murine leukaemia virus-related virus (XMRV), as well as 6,461 VMs, 79,089 VISs (71,595 VISs, sum overlaps > 0), 11,637,065 overlapping entries between VISs and Cistromes, 65 virus-associated diseases (25 public gene expression datasets performing immune analysis), 15,056 target genes and 56 datasets with clinical information. The detailed statistics for ViMIC data are listed in **Table 1**.

**Table 1.** Statistics of ViMIC data.

| **Data Type** | **Count** | **Description** |
| --- | --- | --- |
| Virus | 8 | Including 5 DNA viruses and 3 RNA viruses. |
| Virus Mutation (VM) | 6,461 | Curated mutation information entries. |
| Histone Mark | 78 | Histone Mark types overlapped with VISs in ViMIC. |
| Transcription Regulator | 1,358 | Transcription regulator types overlapped with VISs in ViMIC. |
| Viral Integration Site (VIS) (Sum Overlaps >0) | 71,595 | VISs that overlaps with cistrome data. |
| VIS-Histone Modification (HM) Overlaps >0 | 66,322 | VISs that overlaps with cistrome HM data. |
| VIS-Transcription Factor Binding Site (TFBS) Overlaps >0 | 53,719 | VISs that overlaps with cistrome TFBS data. |
| VIS-Chromatin Accessibility (CA) Overlaps >0 | 29,172 | VISs that overlaps with cistrome CA data. |
| Virus Related Disease | 65 | Disease types which associated with the 8 ViMIC viruses. |
| Target Gene | 15,056 | Curated target genes affected by viral genome insertion or virus gene/protein/region regulation. |
| Literature | 2,153 | Count of the literature related to virus mutation and integration. |
| Clinical Annotation and Data | 56 | Clinical characteristics and virus sequencing data which are available in literatures. |

All of the annotation and analysis results are deposited and managed in a MySQL relational database on a Linux server. Every VM and VIS was assigned unique identifiers, namely, a VVID and DVID, respectively, to perform a cross table query in the SQL database. More detailed information on the VMs, VISs, and target genes is crosslinked to the related resources including PubMed, VISDB, ENCODE, NCBI GenBank (37), Wikipedia, DrugBank (38), NCBI Gene database (39), Human Protein Atlas (HPA), HUGO Gene Nomenclature Committee (HGNC) (40), Vertebrate Genome Annotation (VEGA) (41), Online Mendelian Inheritance in Man (OMIM) (42), STRING (43), Rfam (44), miRbase (45), lncRNASNP2 (46), GEO, TCGA, ESEMBL (47), International Cancer Genome Consortium (ICGC) and European Genome-phenome Archive (EGA) (48).

### Database Utility

#### Data query

ViMIC allows users to search data of interest in three ways. First, a quick search is available on the homepage. Users can input a keyword of interest including a virus name, disease, histone mark, transcription regulator, or virus gene/protein/region. Second, we designed a search engine at the topright corner of each table in the web interface, where users can enter any keywords of interest. Third, an advanced search is also available on the home page for sophisticated searches by selecting a virus name of interest, and searching by human chromosome, disease, virus gene/protein/region or target gene. Each search produces tables of matched entries. Users can then view the detailed information by clicking on the hyperlink or the “View” button in the table.

#### Web interface and data visualization

A web interface was designed to provide user-friendly access to browse, search, analyse, visualize and download data. The database provides an interface for each virus with a summary and links to three ViMIC features to help users quickly check their hypothesis regarding VMs, VIS-cistrome factor overlaps or viral target genes.

First, the “Virus Mutation Site” feature provides the curated mutation information for 8 types of viruses. On the homepage of selected virus, ViMIC presents a catalogue of links to virus genes/ proteins/regions and diseases, and a virus genome plot of each viral reference genome is provided for exploring associations between VMs and genome features. Users can check the location of a particular VM in the selected virus genome together with other well-studied mutations labelled with position, virus gene/protein/region, mutation sites and literature count. Moreover, users can click the “View VM-VIS Association” button to browse the association between reported VMs and reported VISs. The detailed virus mutation page is composed of mutation annotation and evidence sentences from the literatures.

Second, the “Viral Integration Site” feature presents analysis results, including the number of VISs and three cistrome factor-overlapped records. On the selected virus integration page, ViMIC provides the sum of VIS-cistrome factor counts and three VIS-cistrome factors overlap counts in each human chromosome, as well as two browse data menus for users to view overlapped entries by chromosomes or factors. Users can browse VIS-cistrome factor overlapped records data by clicking the “View” button in the table as well as by selecting a chromosome or cistrome factor in the browse data menu. Moreover, ViMIC designs a search module to filter out overlap data according to options of user’s interest like disease, biological source (cell line, cell type and tissue type), etc. In addition, ViMIC provides a heatmap of three VIS-cistrome factors overlap counts on each chromosome and two stacked barplots of top-ranked cistrome factors to help users have a better understanding of the distribution of VIS-cistrome factor overlap in the human genome.

Third, the “Target Gene” feature provides the curated target genes affected by viral genome insertion or by virus gene/region/protein. Users can browse the detailed gene information, related VISs/VMs and the correlation analysis results between gene expression levels and the proportions of 22 types of infiltrating immune cells in virus associated human diseases.

ViMIC also developed virus, disease, dataset, upload, download and help modules at the homepage, which allows users to be quickly redirected to a particular feature of users’ interest. In addition, users can browse datasets for which clinical characteristics and virus sequencing data are available in the literature in our dataset module. To make the data presentation more intuitive, the statistical plots and tables are also shown on the home page of each module.

#### Case exploration

##### Visualization of the virus mutation site

In this article, we take HBV as an example. Users can click the green button with VM number in HBV, as shown in **Figure 2A**. ViMIC will then return a table containing the manually curated information for the HBV mutation sites. Users can quickly enter the keyword in the search box on the top-right of the table, and click the ‘View’ button to explore the detail information of searched mutation, including mutation level, virus gene/protein/region, genotype and evidences mentioned in literatures.

**Figure 2.**
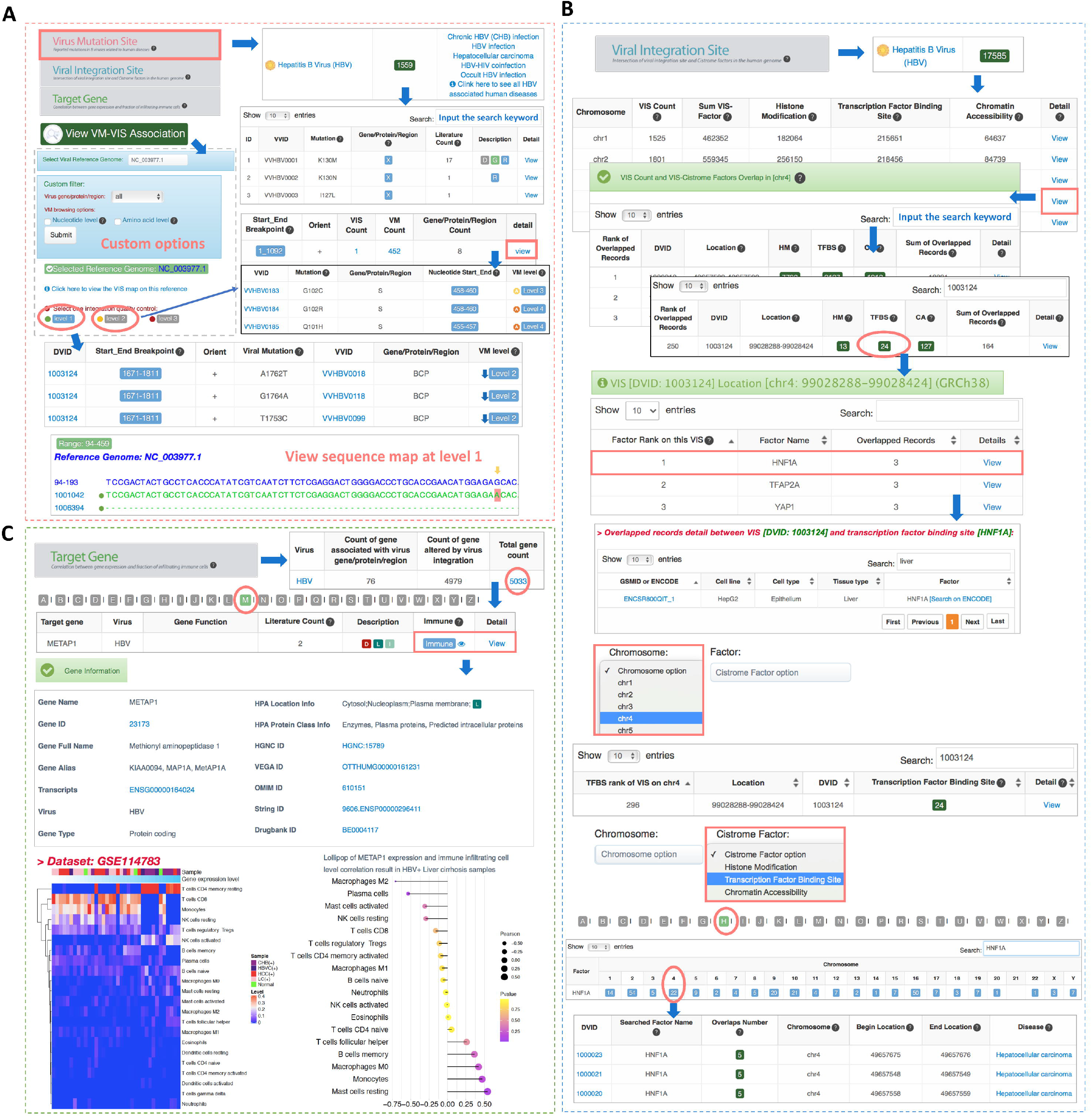
Screenshot depicting an example exploration of the ViMIC database. (A) Assuming the user is interested in HBV, he or she can access a page on ViMIC containing brief mutation information on HBV by clicking HBV VM Count. The user can input a keyword through the quick search engine and enter the detailed mutation information page to view the mutation site. Users can click the “View VM-VIS Association” button to browse the association between VMs and VISs. By selecting a reference genome (e.g., NC_003977.1), ViMIC will return a result showing the distribution of mutations harbored in VISs or the reported VMs in the range of viral integration fragments based on different levels of VIS data. (B) Assuming the user is interested in acquiring a detailed factor list for VIS entry “1003124” on chromosome 4 in HBV, ViMIC will show the overlaps result of three cistrome factors after the users clicks the “chr4” and search the entry of “1003124”. Assuming the user is interested in the “24” overlap number within the “1003124” and “TFBS” entry, ViMIC will then return the ranking of all transcriptional regulators that have overlaps with this VIS. By clicking the ‘View’ button of HNF1A, ViMIC will generate a HNF1A-related table including the GSMID/ENCODE ID, cell line, cell type, tissue type and factor name. On the HBV VIS-cistrome factor homepage, the user can select the chromosome menu to view the overlap distribution of three factors and virus integration fragments on a specific chromosome (e.g., chr4). By selecting the factor menu, the user can explore more overlap statistics and distribution information for a specific transcription factor (e.g., HNF1A). (C) ViMIC provides the “Target Gene” feature for the curated target genes affected by viral genome insertion or by virus gene/protein/region. Assuming the user is interested in the VIS 1003124 target gene METAP1 in HBV, ViMIC will show METAP1 gene information, inserted VISs reported in the literature, and the correlation between gene expression levels and the fraction of infiltrating immune cells in HBV related human disease on the detailed gene information page.

To browse the association of VMs and VISs, users can click “View VM-VIS Association” on the homepage of HBV. ViMIC will show the level 1’s VIS data and matched reported VMs by default. As shown in the example of HBV reference genome NC_003977.1, users can view the distribution of mutations harbored in VISs compared with the NC_003977.1 sequence. The sequence map allows users to see all level 1’s VISs and click DVID to query the detailed information of each VIS, view mutations that have been reported in this range, and enter the VM annotation module by clicking the icon of the VM location. On the other hand, ViMIC designed buttons for users to browse VIS data of other levels (e.g., level 2), if they are interested in which mutations are reported in the range of integration. Additionally, ViMIC set a series of dropdown menu items above the table for users to choose different reference sequence of virus, virus gene/protein/region, and VM browsing options (**Figure 2A)**.

##### Visualization of the viral integration site overlapped with cistrome factors

If users wish to acquire a detailed factor list for one VIS entry, they can follow the step in **Figure 2B**. We take the entry of “1003124” on chromosome 4 in HBV as an example. After users click the “chr4” and search the entry of “1003124”, ViMIC shows the overlaps result of three cistrome factors. By clicking the green button with number at “TFBS” (Transcription factor binding site) column, ViMIC shows the ranking of all transcriptional regulators that have overlaps with this VIS. For instance, users can view cistrome samples by clicking the ‘View’ button of the transcription factor HNF1A (hepatocyte nuclear factor 1 homeobox A), and can find that the sample ENCSR800QIT_1 shows an overlap between “1003124” and HNF1A at the position on chromosome 4 (coordinate range: 99028288-99028424, hg38). ViMIC provides information about this VIS integrating into the host genome which may be a functional element interacting with HNF1A. Moreover, users can also select “chr 4” in the integration menu of HBV to inspect the overlap distribution of three factors and virus integration fragments on human chromosome 4. In addition, users can select the factor browse menu to view all overlapped entries between curated HBV VISs and HNF1A in the human genome (**Figure 2B**).

##### Visualization of target gene information and immune analysis result

As METAP1 is a target gene of “1003124” (49), ViMIC presents the immune analysis provided by CIBERSORT between METAP1 gene expression and 22 types of infiltrating immune cells (**Figure 2C**). On the detailed information page of METAP1, ViMIC shows the heatmap and correlation analysis results for HBV infected diseases as well as providing the METAP1 gene name, ID, full name, alias, transcripts, type, HPA location and protein classification information, and several gene ID links to other resources. For instance, the METAP1 gene expression analysis results show the highest negative correlation with macrophage M2 in HBV+ liver cirrhosis samples.

## DISCUSSION AND FUTURE DIRECTIONS

ViMIC is developed as a comprehensive database of human disease-related viruses from the perspective of VMs, VISs and virus cis-effects, and is a well-maintained resource that will be of great value to different users.

ViMIC collected virus mutation information retrieved from a large collection of published literature. This feature will provide researchers who are interested in VMs with annotations of the reported pathogenic mutations in 8 viruses related to human diseases easily. ViMIC integrated VMs and VISs information from the literatures and it integrates VIS data with cistrome resources. Previous studies pointed out that HBx can enhance virus replication leading to a higher frequency of HBV-DNA integration events into the human genome (11). The HBx sequence easily to integrate into hepatocytes with multiple point mutations, plays an important role in HBV infection and liver cancer development (10). This feature may help spur the further investigation of these two aspects of complicated mechanisms of viral pathogenesis. ViMIC also integrate VIS data with cistrome resources. The viral sequence inserted into the host genome may be a functional element interacting with histone modification, transcriptional regulation and cis-regulatory elements. For example, researcher have found that the transcriptional factor HNF1A contributes to the control of HBV replication (26). The liver-specific activity of one of the critical cis-elements, Enhancer II (nt. 1636-1741, partially contained in VIS 1003124), is regulated by multiple liver-enriched transcription factors, including HNF1A (27). Our work will contribute to and help researchers who are interested in gene regulation mechanisms to systematically investigate the functional annotation of VISs in the host genome. Furthermore, Jungnam Joo et al. (50) indicated that the viral target genes related to immune response may be associated with the viral integration patterns to avoid the immune surveillance system in the body and affect the response to treatment. ViMIC provides three aspects of viral target gene information for users to browse the gene information, VISs insertion information, and the correlation between the expression level of viral target genes and infiltrating immune cell fractions. Additionally, advanced users can conveniently download the annotated virus data and conduct further analysis by themselves.

In the future, ViMIC will be maintained and updated continuously, since VMs, VISs and cistrome data are expected to accumulate. The utility of ViMIC will be improved in several aspects. For example, we will further classify and annotate drug resistance sites and immune escape sites. The sequence around the viral integration site will be downloaded from public databases to perform indepth analysis to investigate the functional role of virus-host interactions. The upload module will be optimized for users to upload potential VISs of interest in batches and perform online analysis. In addition, we will also integrate other immune deconvolution tools such as TIMER (51) and xCell (52), to optimize the anaysis of the association between gene expression and immune cell infiltration in the viral target gene module. In summary, we hope that ViMIC can give the biomedical community a better comprehensive understanding of virus-related research.

## SUPPLEMENTARY DATA

Supplementary Data are available at NAR Online.

## DATA AVAILABILITY

ViMIC is available at http://bmtongji.cn/ViMIC/index.php. The E-Utilities is a set of eight server-side programs that provide a stable interface into the Entrez query and database system available in NCBI (https://www.ncbi.nlm.nih.gov/books/NBK25500/). GIGGLE can search and rank the overlapping genomic loci between query genomic locations such as VISs and genome interval files and it is available as a GitHub repository at https://github.com/ryanlayer/giggle. LiftOver is a web service that provides genome coordinate transformation between different human genome reference assemblies and can be freely accessed at https://genome.ucsc.edu/cgi-bin/hgLiftOver. PubTator is a web service that can automatically annotate biomedical entries including genes and mutations, from the literature, which is available at https://www.ncbi.nlm.nih.gov/research/pubtator/. CIBERSORT uses the gene expression data to estimate the abundances of member cell types from a mixed cell population, and it can be freely accessed at https://cibersort.stanford.edu/.

## ACKNOWLEDGEMENTS

We are grateful to Dr. Shipeng Chen for reading the manuscript.

## FUNDING

This work was supported by the National Natural Science Foundation of China [81972914, 81573023], the Fundamental Research Funds for the Central Universities [22120200014], Shanghai “Rising Stars of Medical Talent” Youth Development Program [2019-72], and the National Key R&D Program of China [2016YFC1303200].

## CONFLICT OF INTEREST

The authors disclose no conflicts of interest.

## Notes

### Competing Interest Statement

The authors have declared no competing interest.

